# In vitro evaluation of the effect of mutations in primer binding sites on detection of SARS-CoV-2 by RT-qPCR

**DOI:** 10.1101/2021.06.07.447338

**Authors:** Fee Zimmermann, Maria Urban, Christian Krüger, Mathias Walter, Roman Wölfel, Katrin Zwirglmaier

**Affiliations:** Bundeswehr Institute of Microbiology, Neuherbergstr, 11, 80937 Munich, Germany

**Keywords:** SARS-CoV-2, RT-PCR, primer mismatch, diagnostics

## Abstract

A number of RT-qPCR assays for the detection of SARS-CoV-2 have been published and are listed by the WHO as recommended assays. Furthermore, numerous commercial assays with undisclosed primer and probe sequences are on the market. As the SARS-CoV-2 pandemic progresses, the virus accrues mutations, which in some cases – as seen with the B.1.1.7 variant – can outperform and push back other strains of SARS-CoV-2. If mutations occur in primer or probe binding sites, this can impact RT-qPCR results and impede SARS-CoV-2 diagnostics. Here we tested the effect of primer mismatches on RT-qPCR performance in vitro using synthetic mismatch in vitro transcripts. The effects of the mismatches ranged from a shift in ct values from −0.13 to +7.61. Crucially, we found that a mismatch in the forward primer has a more detrimental effect for PCR performance than a mismatch in the reverse primer. Furthermore, we compared the performance of the original Charité RdRP primer set, which has several ambiguities, with a primer version without ambiguities and found that without ambiguities the ct values are ca. 3 ct lower. Finally, we investigated the shift in ct values observed with the Seegene Allplex kit with the B.1.1.7 SARS-CoV-2 variant and found a three-nucleotide mismatch in the forward primer of the N target.

## 1. Introduction

As part of the SARS-CoV-2 response the international scientific community rapidly published a number of RT-PCR diagnostic assays for SARS-CoV-2 detection. 15 RT-PCR assays of national reference laboratories are listed by the WHO for SARS-CoV-2 detection in patient samples (WHO, 2020). Although SARS-CoV-2 has a slower mutation rate than, for example, influenza (Manzanares-Meza and Medina-Contreras, 2020), it nevertheless has accrued mutations in most of the primer and probe binding sites of globally used PCR assays within a few months of the start of the pandemic. Aligning 177 genomes collected up to July 2020 from across Brazil with primer sequences of 15 of the WHO-listed SARS-CoV-2 assays revealed that only 3 assays (NIID_2019-nCoV-N, nCoV-IP4 and CN-CDC-E) had a perfect match to all 177 genomes (Santos et al., 2020). A similar study using sequence data from 375 genomes collected around the world up to April 2020 showed that only 2 out of 12 WHO-listed assays had a perfect match to all analyzed genomes (CDC-N1 and Charité RdRP) (Toms et al., 2020). Depending on location and nature of the mismatches this could have significant effects on assay performance and negatively impact sensitivity. The possible effect of primer mismatches in silico has been examined in a number of publications (Bru et al., 2008; Huang et al., 1992; Kwok et al., 1990; Lefever et al., 2013; Stadhouders et al., 2010). However, in vitro results don’t always agree with in silico analyses, since various confounding factors can influence the outcome (Lefever et al., 2013; Mendelman et al., 1990; Stadhouders et al., 2010). Therefore, the aim of this study was to check the impact of in silico described primer mismatches on the in vitro performance of several of the WHO-listed SARS-CoV-2 RT-PCR assays. Furthermore, approaching the issue from the other side, we wanted to investigate the in silico basis of an in vitro observed dropout of a PCR target of a commercial SARS-CoV-2 assay.

## 2. Material and Methods

### 2.1. SARS-CoV-2 strains used in this study

We used extracted RNA from SARS-CoV-2 strain mucIMB-1 to generate the in vitro transcripts of the mismatch and perfect match templates. mucIMB-1 (EPI_ISL_406862) was isolated in January 2020 from a patient in Bavaria, Germany, and is phylogenetically close to the first published SARS-CoV-2 sequence, Wuhan-Hu1. For investigating the performance of several commercial SARS-CoV-2 PCR kits with SARS-CoV-2 variant B.1.1.7, we used strain mucIMB-CB (EPI_ISL-755639) in comparison with mucIMB-1. mucIMB-CB was isolated from a patient with a history of travel from the UK in December 2020.

### 2.2. Selection of primer mismatches

The European Centre for Disease Control (ECDC) provides an online tool “PrimerScan” that regularly screens all available SARS-CoV-2 genomes published on GISAID for mutations in the primer and probe binding sites of WHO recommended RT-PCR assays (https://primerscan.ecdc.europa.eu/). The website also plots the frequency of these mutations over time. Based on this information we chose several of the most frequent mutations between March and September 2020 for this study. Several of these mutations have since disappeared again.

### 2.3. Construction of mismatch templates

In order to generate RNA templates with the required mismatches in the primer binding sites, we used primers containing the mismatch (or complement base of the mismatch for reverse primers) (Supplementary table 1) and performed an RT-PCR with SARS-CoV-2 mucIMB-1 strain (which has no mismatch to any of the PCR assays tested here). For several mismatches we had to relax the annealing temperature in order to be able to generate a PCR product (Table 1). Both forward and reverse primers contained a short nonsense sequence 5′ of the primer sequence so as to generate a PCR product that is slightly longer than the template for the subsequent experiments. The purpose is to avoid artefacts due to primers binding right at the end of a template. The forward primers carried the T7 promotor at the 5′ end. We then used in vitro transcription to produce the mismatch templates. Correct (i.e. no mismatch) templates were produced in the same way.

**Tab. 1.** Effect of mismatches in template on PCR. * RdRP primer RdRP_SARSr-F2/RdRP_SARSr-R1.corrECDC (with ambiguities), § RdRP primers RdRP_SARSr-F2.new/RdRP_SARSr-R1.new (without ambiguities)

| Mismatch | max frequency % /date | Mutation | Mismatch type | distance from 3' end | delta ct compared to wildtype | Annealing temp used to generate mismatch template |  |
| --- | --- | --- | --- | --- | --- | --- | --- |
|  | data from ECDC PrimerScan |  | primer : mismatch IVT |  |  | normal PCR | to generate mismatch template |
| RdRP MM F8 | 0.46 /06-2020 | G => T | G : A | 15 | -0.01*/+0.16 § | 58 | 58 |
| N_Sarbeco MM R8 | 1.37 /06-2020 | C => A | G : A | 13 | 3.48 | 58 | 56 |
| N_Sarbeco MM R13 | 2.40 /06-2020 | C => A | G : A | 8 | 1.05 | 58 | 56 |
| CDC N1 MM R15 | 0.17 /03-2020 | C => T | G : T | 10 | 6.79 | 55 | 55 |
| CDC N1 MM R21 | 0.50 /08-2020 | T => G | A : G | 4 | 2.31 | 55 | 55 |
| CDC N1 MM R22 | n.d. | G => T | C : T | 3 | 1.54 | 55 | 55 |
| CDC N3 MM F8 | 4.18 /02-2020 | T => C | T : G | 15 | 4.37 | 55 | 55 |
| CDC N3 MM R14 | 0.17 /02-2020 | G => T | C : T | 8 | 2.86 | 55 | 55 |
| CDC N3 MM F10 | 0.39 /06-2020 | G => T | C : T | 13 | 1.6 | 55 | 55 |
| Japan N MM F20 | 0.27 /07-2020 | C => T | C : T | 1 | 7.61 | 60 | 60 |
| Japan N-neu MM R17 | 0.50 /05-2020 | G => C | C : C | 4 | 3.44 | 60 | 51 |
| Japan N-neu MM R18 | 0.64 /01-2020 | T => G | A : G | 3 | 5.25 | 60 | 51 |
| IP2 MM R8 | n.d. | A => G | T : G | 11 | -0.13 | 58 | 56 |
| IP2 MM R17 | 0.45 /06-2020 | C => T | G : T | 2 | 2.3 | 58 | 58 |

The detailed protocol is described below.

To generate the DNA templates needed for in vitro transcription PCR was performed on a Thermo™ Scientific Piko™ Thermal Cycler (Thermo Fisher Scientific, Waltham, MA, USA) using the QIAGEN One-Step RT-PCR Kit (QIAGEN, Hilden, Germany) in 50 μl reactions containing 10 μl 5x QIAGEN OneStep RT-PCR Buffer, 1.4 μl dNTP Mix (10 mM of each dNTP), 0.6 μl dNTP Mix dUTP Solution (premix of 10 mM dATP, dCTP, dGTP and 20 mM dUTP; Jena Bioscience GmbH, Jena, Germany), 1.5 μl Antarctic UDG (New England Biolabs, Ipswich, MA, USA) and 2 μl QIAGEN OneStep RT-PCR Enzyme Mix. 5 μl (2 x 10e4 copies/μl) SARS-CoV-2 RNA were used as template. The amplification conditions were 50 °C for 30 min, 95°C for 15 min, followed by 45 cycles of 94 °C, 15 sec and assay specific annealing temperature according to Table 2 for 30 sec and a final elongation at 72 °C for 30 sec. PCR products were analyzed by 2 % agarose gel electrophoresis. Bands corresponding to the correct fragment size were cut out and the DNA extracted using the QIAquick Gel Extraction Kit (QIAGEN, Hilden, Germany). The sequence was confirmed by Sanger sequencing.

**Table 2.**
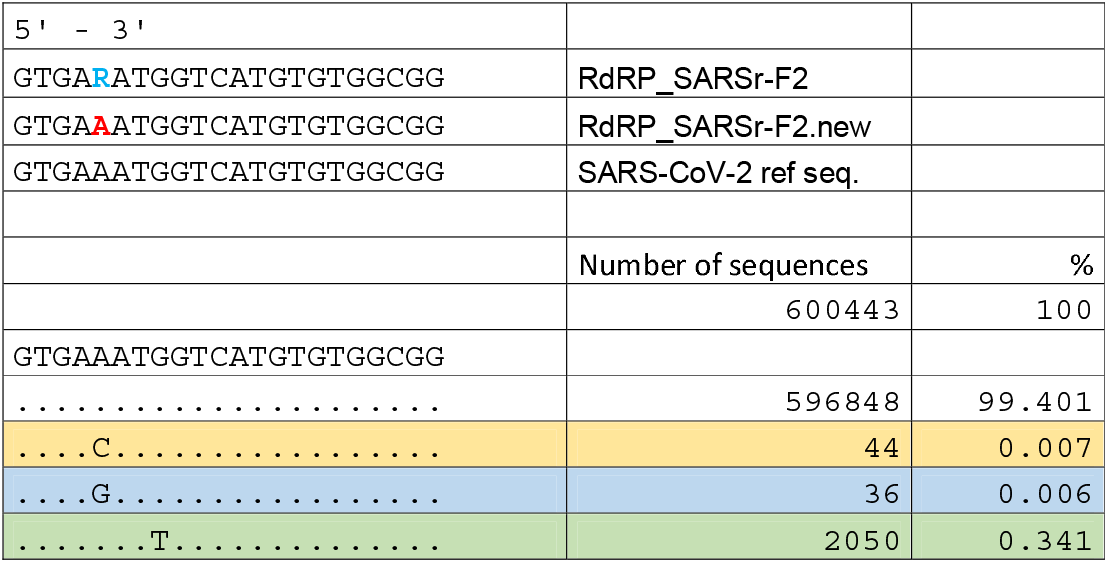
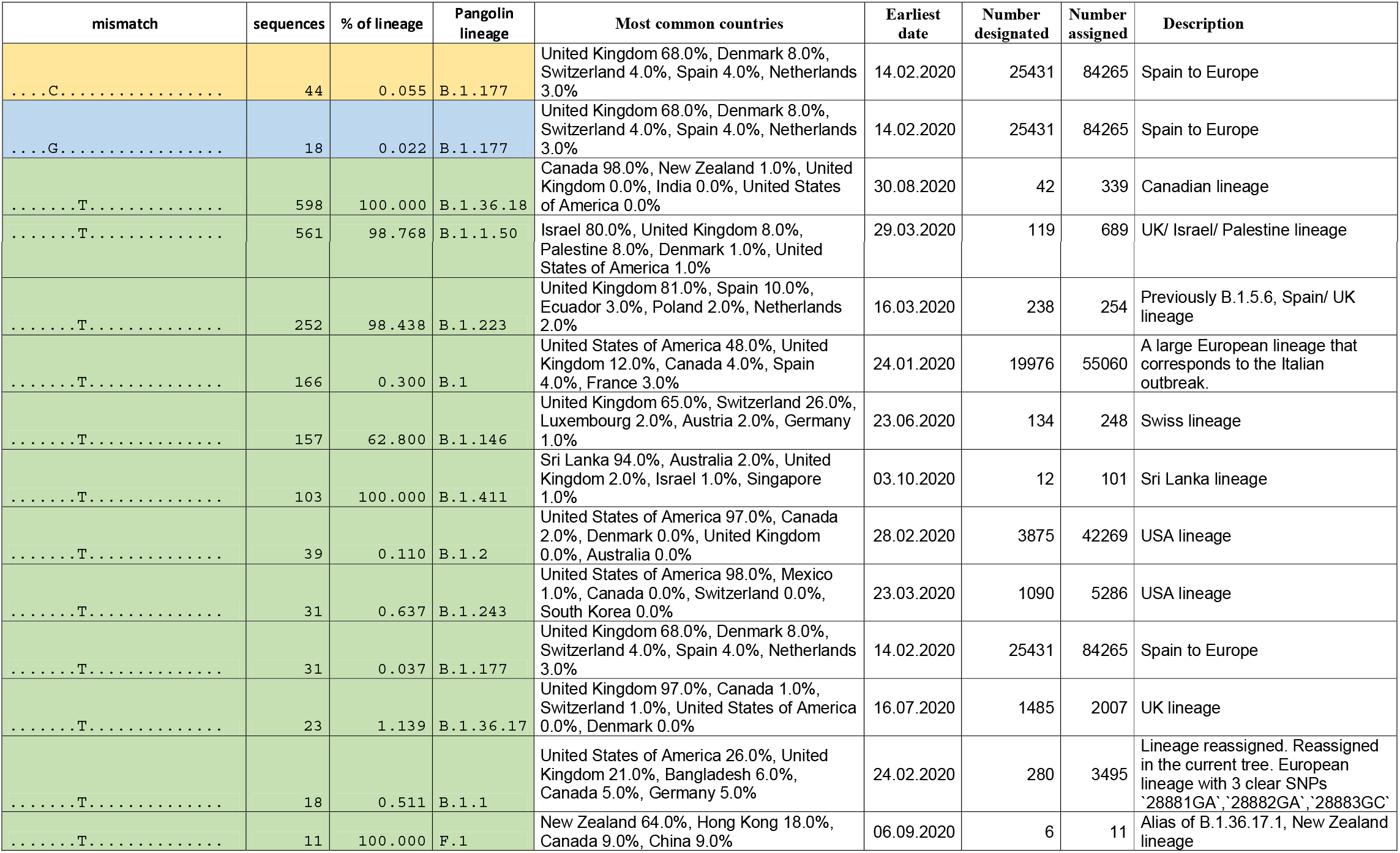
Mismatches of Charité RdRP forward primer in 600 000 published SARS-CoV-2 genomes. Table 2a. Number of sequences with mismatches in specific positions Table 2b. Lineage analysis of the mismatches. Lineage information retrieved from Pangolin lineages (https://cov-lineages.org/) on 15.03.2021. Only lineages where > 10 sequences carry the mutation are shown

RNA templates were produced by in vitro transcription using the MEGAscript T7 Kit (Thermo Fisher Scientific, Waltham, MA, USA) following the manufacturer’s protocol including a TURBO DNase treatment step (15 min at 37°C). The in vitro transcripts (IVTs) were cleaned up using the NucleoSpin RNA Clean-Up Kit (Macherey-Nagel, Düren, Germany) according to manufacturer’s instructions and subsequently quantified by using a Qubit RNA BR Assay Kit (Thermo Fisher Scientific, Waltham, MA, USA). The copy number was calculated using the online NEBio Calculator (https://nebiocalculator.neb.com/#!/ssrnaamt).

### 2.4. RT-PCR with correct and mismatch IVTs

Dilution series (triplicates) of in vitro transcripts of the correct and mismatch templates were amplified by RT-qPCR using the published primers and following the published protocols (ref WHO and references in Supplementary Table 1). All experiments were done with Qiagen OneStep RT-PCR reagents (Qiagen, Hilden, Germany) on a MIC thermocycler (BMS, Brisbane, Australia).

### 2.5. Detecting SARS-CoV-2 variant B.1.1.7 with several commercial PCR kits

Extracted RNA of strains mucIMB-1 (wild type SARS-CoV-2) and mucIMB-CB (B.1.1.7 variant) was used with i) the Seegene Allplex SARS-CoV-2/FluA/FluB/RSV assay (Seegene, Seoul, Korea) and run on a Biorad CFX96DX cycler (Hercules, CA, USA), ii) the VitaPCR SARS-CoV-2 assay (Credo Diagnostics, Singapore) and iii) the GeneXpert Xpress SARS-CoV-2 assay (Cepheid, Sunnyvale, CA, USA). Each assay was performed according to the respective manufacturer’s instructions.

### 2.6. Datamining

We downloaded a total of 600,443 sequences and their corresponding metadata from GISAID at 2021-02-24 10:05. Then we used the search_oligodb command from usearch (Edgar, 2010) to search for the non-degenerated primer sequences (see tables 2a, 3a, 4a) in every SARS-CoV-2 genome sequence with a maximum divergence of three (parameter: -maxdiffs 3). The result sets were enriched with the corresponding pangolin lineage (Rambaut et al., 2020) obtained from the metadata.tsv and statistics were generated using csvtk (Shen, 2021). For some strains, no pangolin lineage was provided with the metadata.tsv. In such case, we classified the strains using pangolin client v2.3 (O’Toole et al., 2021).

**Table 3.**
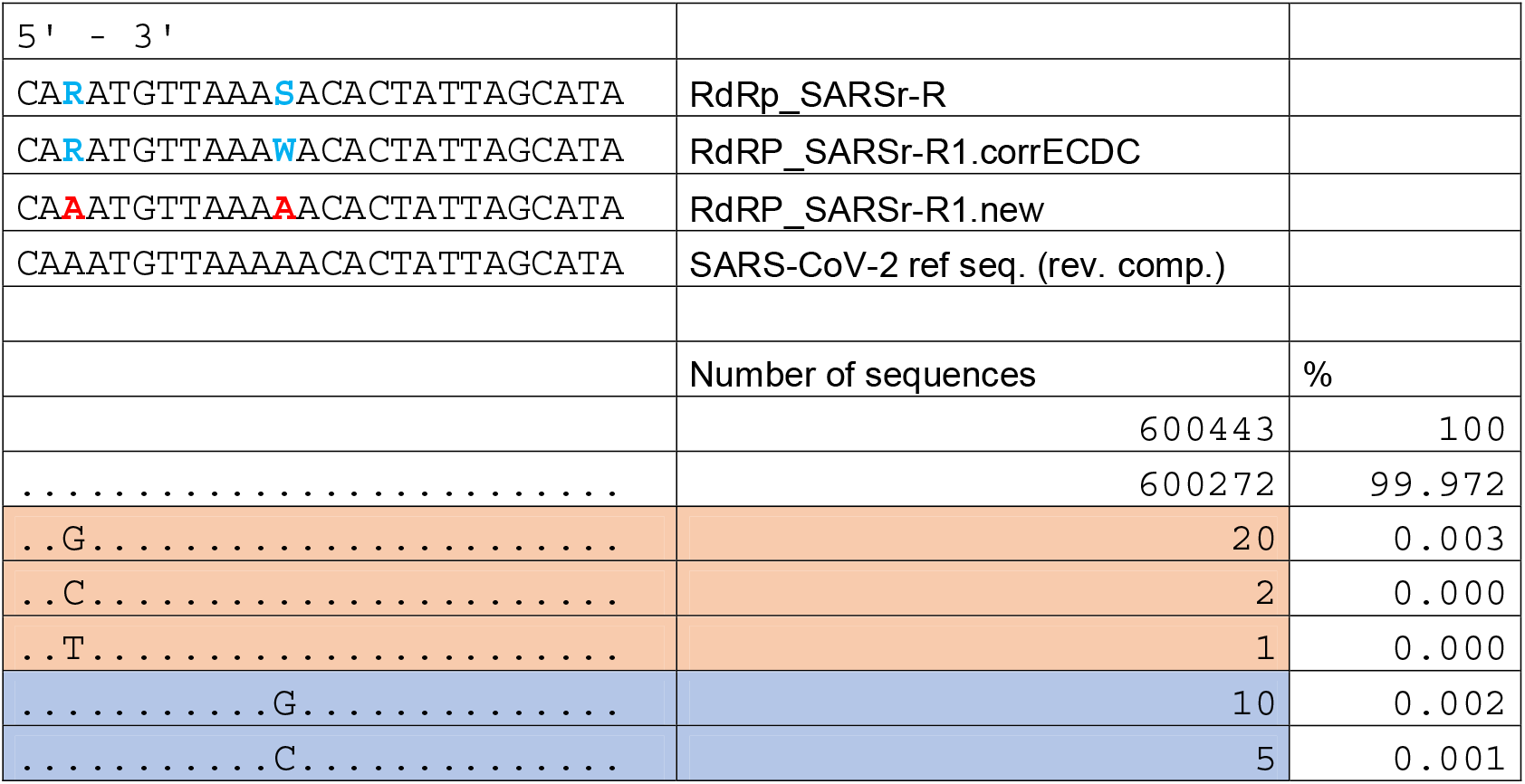
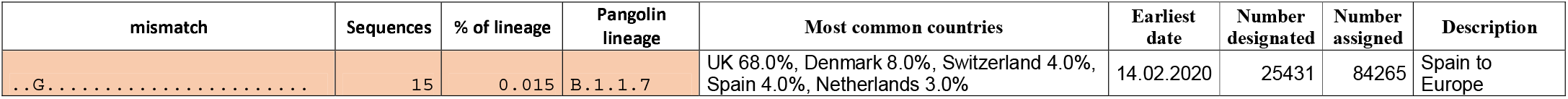
Mismatches of Charité RdRP reverse primer in 600 000 published SARS-CoV-2 genomes. Table 3a. Number of sequences with mismatches in specific positions Table 3b. Lineage analysis of the mismatches. Lineage information retrieved from Pangolin lineages (https://cov-lineages.org/) on 15.03.2021. Only lineages where > 3 sequences carry the mutation are shown

**Table 4.**
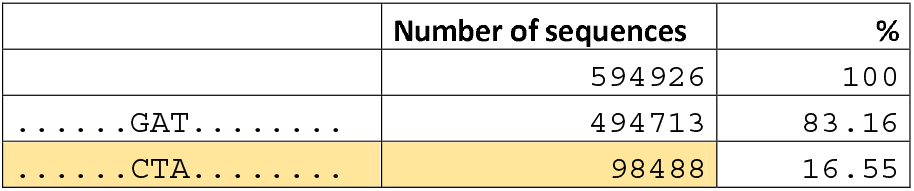
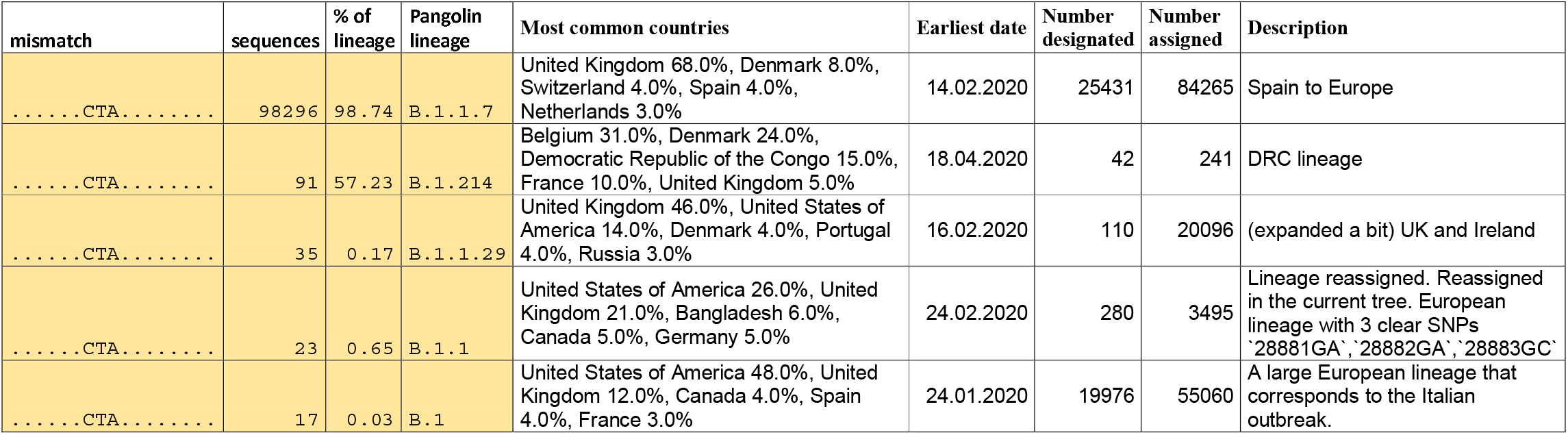
Triple mismatch in the Seegene Allplex N-assay in genome position 28280-28282. Table 4a. Number of sequences with the triple mismatch Table 4b. Lineage analysis of the mismatches. Lineage information retrieved from Pangolin lineages (https://cov-lineages.org/) on 15.03.2021. Only lineages where > 10 sequences carry the mutation are shown

## 3. Results

### 3.1. Mismatches in WHO-listed primers

Based on the ECDC PrimerScan tool (https://primerscan.ecdc.europa.eu) we chose 14 mismatches in primer binding sites of six different WHO-listed RT-qPCR assays. We generated in vitro transcripts of the mismatch templates and then performed RT-qPCR with the published primers, comparing the ct values obtained with the “correct” templates and the “mismatch” templates (Table 1 and Figure 1). The effect of the mismatch on the ct values ranged from −0.13 to +7.61. Within a dilution series ranging from 10e5 copies/reaction to 10e0 copies/reaction the observed shift in ct values between the correct and the mismatch template was consistent across all dilutions, i.e. independent of the template concentration. Regarding the distance of the mismatch from the 3′ end of the primer and the magnitude of the ct shift, there was only a weak correlation (Pearson −0.320) for the entire dataset. However, when analyzing forward and reverse primer separately, we found a strong negative correlation for mismatches in the forward primer (Pearson −0.822) and no correlation for the reverse primer (Pearson 0.004). Analyzing the data from a recent study on primer mismatches in SARS-CoV-2 RT-qPCR assays (Nakabayashi et al., 2021) under this aspect, we found the same trend, i.e. a strong negative correlation between the distance of the mismatch from the 3′ end and the magnitude of the ct shift for the forward primer (Pearson −0.576) and only a weak correlation for the reverse primer (Pearson −0.159).

**Fig. 1.**
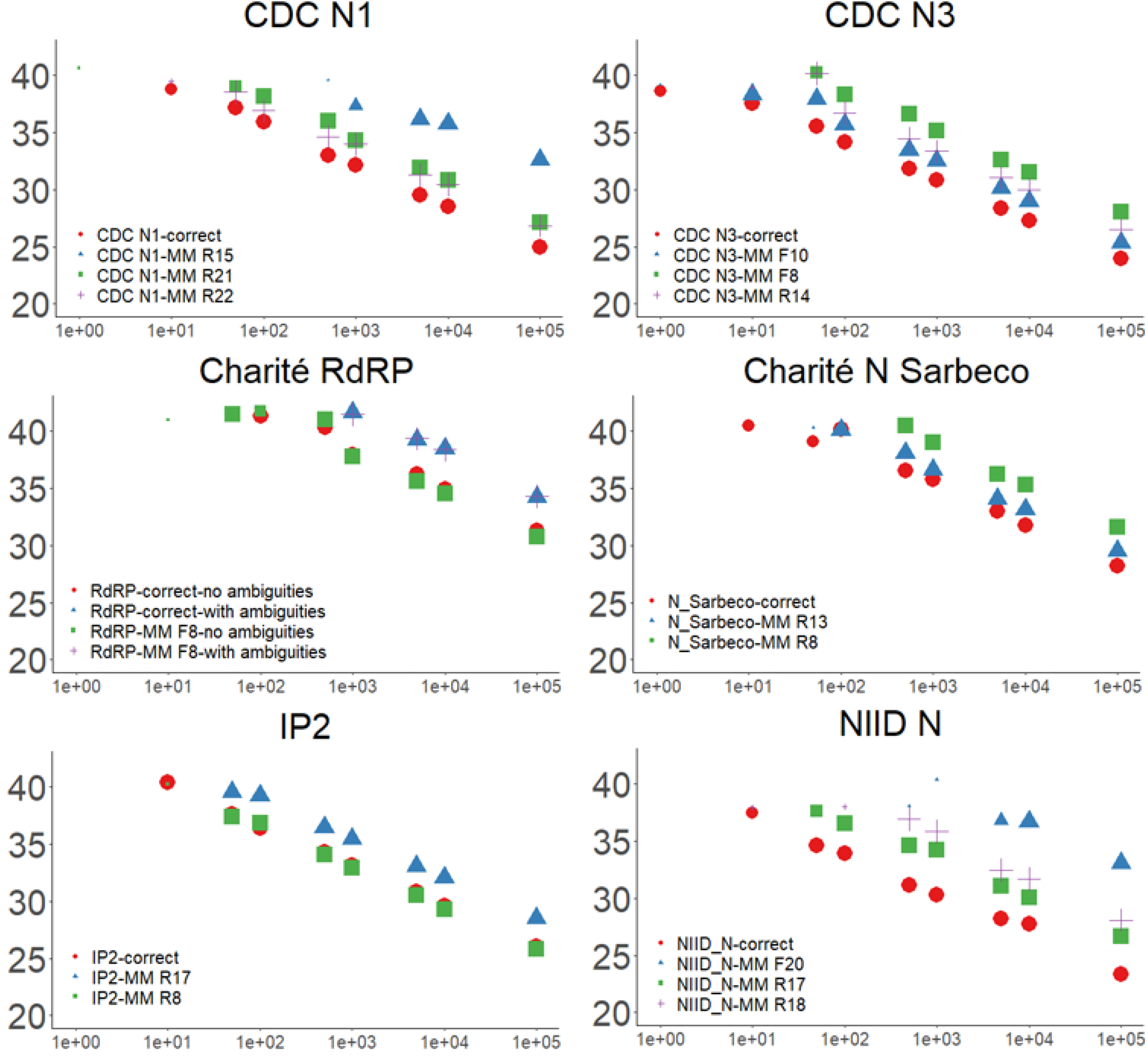
PCR results using the published primers with the correct templates and mismatched templates. Dilution series were done in triplicate. Size of the symbols represents one (small), two (medium sized) or three (large) positive replicates.

With regard to the mismatch type, there was no consistent pattern. A G:T mismatch is thought to be generally well tolerated in PCR (Kwok et al., 1990; Rejali et al., 2018; Stadhouders et al., 2010). However, in our study a G:T mismatch resulted in a large delta ct of 6.79 at a distance of 10 bases from the 3′ end (CDC-N1-MM R15) in one instance and a delta ct of only 2.3 at a distance of 2 bases from the 3′ end (IP2-MM R17) in another instance.

### 3.2. Charité RdRP primer set

The original Charité primer set for RdRP (Corman et al., 2020) carries three ambiguous positions, one in the forward primer (position 5) and two in the reverse primer (positions 3 and 12 (Supplementary table 1). We conducted an in silico analysis of the primer binding sites using > 600,000 SARS-CoV-2 genomes published on GISAID (as of February 2021). 99.40 %of genomes carry an A at position 5 in the forward primer and 99.97 % carry a T (i.e. an A in the primer) at both positions 3 and 12 of the reverse primer (Tables 2a and 3a). We therefore used primers with the dominant sequence (designated RdRP-SARSr-F2.new and RdRP-SARSr.R1.new) and compared their performance to primers containing the ambiguities. Both the correct template as well as a template including a mismatch in the binding site of the forward primer, RdRP MM F8, yielded PCR products with lower ct values (−3.16 for correct template, −3.60 for MM F8) with the .new primers compared to the original Charité primers (Fig. 1a). In both primer sets (Charité and .new) the difference in ct values between the correct template and mismatch template was negligible (Table 1), indicating that the mismatch has no pronounced effect on amplification.

Further in silico analysis showed that for the reverse primer sequence only 23 respectively 15 out of 600,443 SARS-CoV-2 genomes carry a mismatch to the .new primer sequence in positions 3 respectively 12. The mismatches are spread over eight (position 3) and nine (position 12) lineages with no clear cluster in any one lineage (Table 3b). No genome carries both mismatches at the same time. For the forward primer 44 genomes carry a C and 36 a G at position 5 of the primer. The C mismatch only appears in lineage B.1.177 (0.052 % of B.1.177 genomes), while the G mismatch is spread over 9 lineages. A more dominant mismatch is the mismatch in position 8 of the forward primer, mentioned above. This mismatch was investigated in vitro, as 2,050 genomes carried a T, instead of a G (0.34 % of all genomes). This appears in 41 different lineages and is dominant (i.e. present in > 98 % of genomes) in 5 lineages (Table 2b).

### 3.3. SARS-CoV-2 variant B.1.1.7 amplified with several commercial PCR assays

We amplified RNA from wild type SARS-CoV-2 and variant B.1.1.7 with the GeneXpert Xpress SARS-CoV-2 assay, Vita PCR SARS-CoV-2 assay and Seegene Allplex SARS-CoV-2/FluA/FLuB/RSV assay. According to the manufacturers, GeneXpert targets the N and E gene, VitaPCR has two N gene targets and Seegene has targets in the RdRP, S and N gene. Neither GeneXpert nor VitaPCR showed any difference in ct values between wild type and B.1.1.7 (data not shown). The Seegene assay, however, showed a drop in ct values of 7.2 ct in the N target, but not in the S or RdRP target. The drop in ct values was consistent across template concentrations between 10 and 10e5 copies/reaction and led to a loss of sensitivity of 2 log for the N target (Fig 2). We cloned and sequenced the PCR product to identify the mismatch causing the drop in PCR performance. Aligning the PCR amplicon to SARS-CoV-2 WuhanHu1 and B.1.1.7 revealed a triple mismatch in the forward primer (GAT => CTA), corresponding to SARS-CoV-2 genome positions 28,280-28,282 at the start of the N gene (Fig. 3). This mismatch currently (as of February 2021) has an overall frequency of 16.55 % in 600,000 genomes on gisaid.org and appears in 98.74 % of all B.1.1.7 genomes (UK variant) and 57.23 % of all B.1.214 genomes (DRC variant) as well as three further lineages with < 1 % frequency (Table 4).

**Fig. 2.**
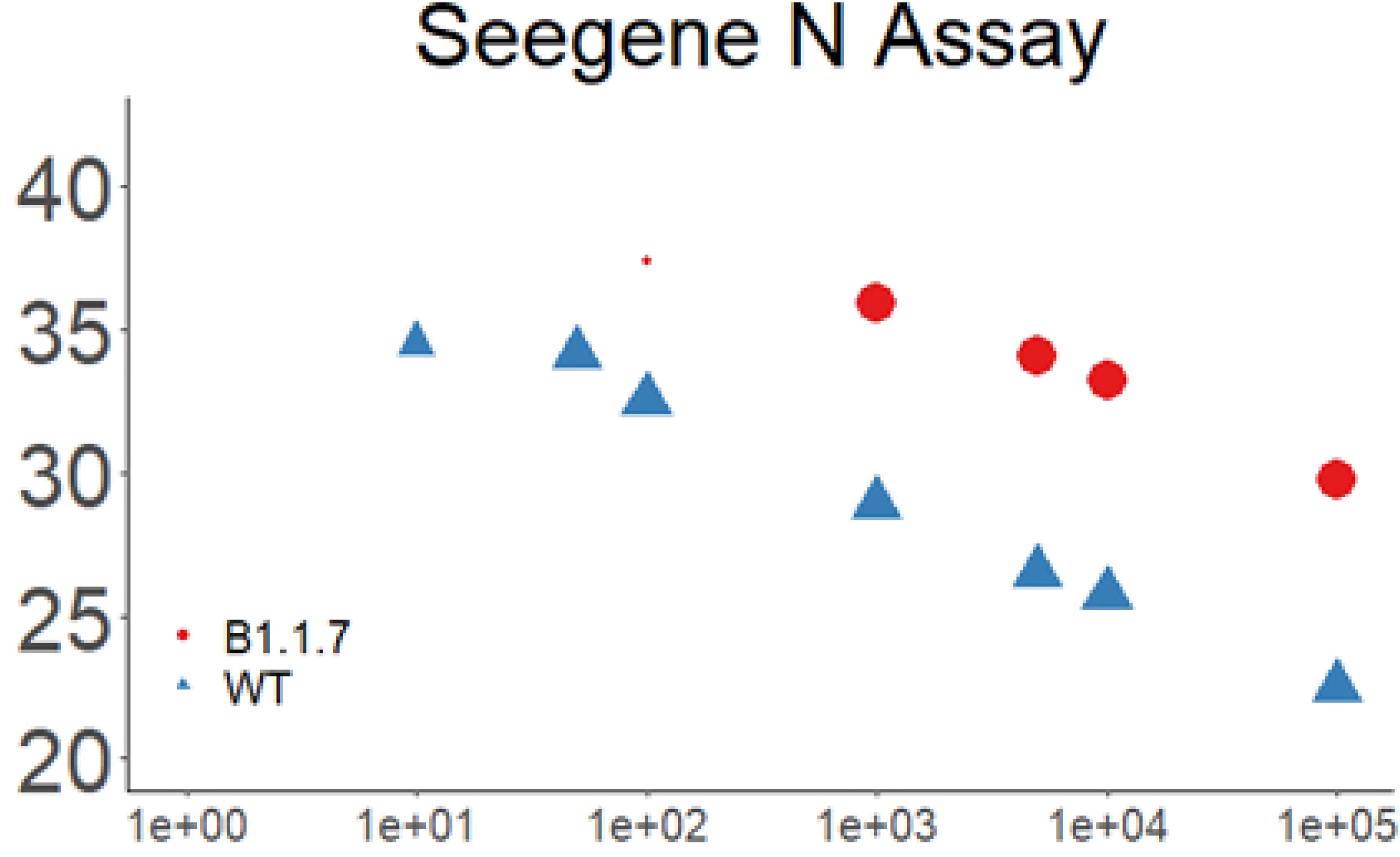
PCR results using the Seegene Allplex assay with SARS-CoV-2 wild type and variant B.1.1.7. Dilution series were done in triplicate. Size of the symbols represents one (small), two (medium sized) or three (large) positive replicates.

**Fig. 3.**
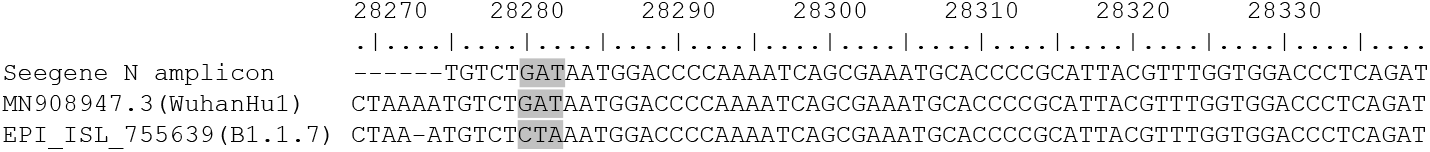
Alignment of the 5′ end of the amplicon of the N-assay of the Seegene Allplex against WuhanHu1 and B1.1.7

## 4. Discussion

### 4.1. Effect of primer mismatches

Several factors can influence the impact of a mismatch on PCR amplification. One intuitive and well documented factor is the proximity of the mismatch to the 3′ end of the primer (Bru et al., 2008; Kwok et al., 1990; Stadhouders et al., 2010). Interestingly, in our study this correlation was not obvious when looking at the whole dataset, but was only true for mismatches in the forward primers. SARS-CoV-2 is a single-stranded positive-sense RNA virus, which means that in an RT-PCR reaction, the first step, the reverse transcription, involves the binding of the reverse primer to the sense RNA strand. The forward primer is only used during the subsequent PCR step. Studies from the early days of PCR noted that reverse transcriptases are more tolerant towards mismatches than DNA polymerases (Huang et al., 1992; Mendelman et al., 1990). Therefore, in an RT-PCR reaction, the reverse transcriptase is more likely to be able to create a cDNA strand from a mismatched primer, correcting the mismatch in the process, while the Taq polymerase is more discriminant against mismatches, resulting in decreased amplification. Thus, in the case of SARS-CoV-2, a mismatch in the forward primer should have a stronger negative impact than in the reverse primer. Our data confirms this, albeit with a limited number of data points. For SARS-CoV-2 RT-PCR assays, this hypothesis is also supported by data from Nakabayashi et al. (Nakabayashi et al., 2021). In an earlier study using HIV-RT-PCR assays, Stadhouders et al. (Stadhouders et al., 2010) made a similar observation. They also noted that when using rTth polymerase (an enzyme that has both reverse transcription and DNA polymerase functionality) instead of a combination of MMLV reverse transcriptase and Taq polymerase for the RT-PCR reaction, mismatches in the forward primer had almost no effect, but mismatches in the reverse primer had a detrimental effect. The authors hypothesize that the different incubation temperatures of MMLV and rTth of 60 °C and 48 °C as well as differences in ionic strength of the reaction buffers contribute to this phenomenon. The varying influence of different PCR reagents on mismatches was also observed by Levefer et al. (Lefever et al., 2013).

Another factor impacting PCR amplification is the mismatch type. A transversion, i.e. resulting in a mispairing of two pyrimidines (C:C, T:T, C:T, T:C) or two purines (A:A, G:G, A:G, GA) is thought to have a greater impact than a transition, i.e. resulting in a mispairing of a pyrimidine and a purine (A:C, C:A, T:G, G:T), mainly due to sterical reasons (Huang et al., 1992; Rejali et al., 2018; Stadhouders et al., 2010). This trend is not obvious in our dataset, particularly illustrated by the example of a G:T mismatch, which had a small effect in close proximity to the 3′ end but a rather large effect further away. A recent study on primer mismatches in SARS-CoV-2 RT-PCR assays has shown the effect from a transition mismatch type in the ultimate 3′ position of a primer to result in a shift in ct values ranging from 0.51 to 6.29 up to a complete PCR failure (Nakabayashi et al., 2021). Taken together, this indicates that not only the mismatch type and position, but also the surrounding sequence of the template and primer influence the outcome, presumably due to secondary structures and interactions based on these (Mendelman et al., 1990; Rejali et al., 2018). Consequently, this serves as a warning that in silico analysis of the effect of primer mismatches can only give an estimate on the impact on PCR performance and should always be tested in vitro.

### 4.2. RdRP primers

The Charité RdRP assay (Corman et al., 2020) was published on 23^th^ Jan 2020, when only the original Wuhan-Hu1 sequence of SARS-CoV-2 was available as a reference. Based on alignments with human SARS-CoV-1 and bat-related SARS-CoV several ambiguities were introduced in the primer sequences. Particularly the base at position 12 from the 5′ end of the reverse primer has sparked some discussion since. Corman et al. inserted an S (= C or G) at this position. Pillonel (Pillonel et al., 2020), after noticing a poorer performance of the RdRP assay compared to the E assay and comparing 1623 SARS-CoV-2 genome sequences available by May 2020, suggested using an R (= A or G), while the ECDC PrimerScan website lists an ECDC-corrected version of the primer with a W (= A or T). Vogels (Vogels, 2020) also noted the poor performance of the RdRP assay and stated that 990 of 992 genomes available at the time carried a T at position 12 of the reverse primer. More than a year into the pandemic and with over 600,000 SARS-CoV-2 genomes sequenced it turns out that 99.97 % carry a T at position 12 as well as position 3, where the Charité reverse primer has the second ambiguity. Since the use of ambiguous primer sequences essentially reduces the amount of “correct” primer for the template (with two ambiguities in the reverse primer, R and S, this results in four primer variants, i.e. only 25 % of the reverse primers in the PCR reaction have the correct sequence) this would account for the poor performance of the primer. Using the primer without ambiguities, with A (to complement the T on the template) in positions 3 and 12 improves the sensitivity of the assay by about 1 log.

For the forward primer the situation is more complex. Only a minute fraction of genomes (0.01 %) carry a G instead of an A at position 5, which would be covered by the ambiguity in the Charité primer sequence. However, we discovered another mismatch in position 8 which, although only present in 0.34 % of the total genome dataset, is the dominant variant in several smaller lineages. This highlights the problem that primer mismatches in localized outbreaks may go unnoticed, while still causing problems with diagnostic assays locally.

### 4.3. Commercial SARS-CoV-2 RT-PCR assays

The 7.2 ct loss in the N target of the Seegene Allplex assay we observed with the B.1.1.7 variant exemplifies the problem with commercial SARS-CoV-2 PCR assays. With proprietary primer and probe sequences the users cannot (or only with considerable effort) check for themselves whether new SARS-CoV-2 variants affect the performance of the assay. They have to rely on communication of manufacturers or serendipitous findings. Other examples of PCR failures with commercial assays were described by Ziegler et al (Ziegler et al., 2020), who found a drop out of the N target in GeneXpert (Cepheid) in one patient sample. Analysis of the sequence revealed a point mutation in the N gene and comparison with GISAID data showed a frequency of 0.2 % for that SNP. Furthermore, Artesi et al. (Artesi et al., 2020) noted a drop out of the E gene target in the Cobas system (Roche) in 0.2 % of SARS-CoV-2 positive samples from Belgian healthcare workers in March/April 2020. The identified SNP at position 26,340 in the E gene was present in just 0.09 % of global SARS-CoV-2 genomes on GISAID. This reiterates the problem pointed out in the last paragraph – not all mutations are as successful as B.1.1.7 and many appear only within a limited geographical region. Another example is a cluster of SNPs in the N gene found in Madeira County, California, USA, that resulted in a loss of 5.4 ct in the Japan N assay NIID_2019-nCoV (Vanaerschot et al., 2021). These regionally restricted mutations may also disappear over time, but while they exist, they can impede diagnostics. This is particularly problematic in countries that do not have the capacity to retest samples with multiple assays or follow up suspicious samples with whole genome sequencing.

On the other hand, a drop out of a PCR target can also be used to screen for a certain mutation, provided that at least a second target is used to confirm SARS-CoV-2. An example is the drop out of the S gene target in the Thermo Fisher TaqPath assay, which is widely used across the UK. It guided and supported the genome sequencing effort that documented the fast spread of variant B.1.1.7 in Britain (Chand et al., 2020).

### 4.5. Conclusions

This in vitro study provides data for the effect of primer mismatches on PCR performance. It highlights the fact that experimental data does not necessarily follow the theoretical predictions, particularly with regard to the magnitude of the ct shift with mismatches close to the 3′ end. This emphasizes the importance of using more than one target in a diagnostic SARS-CoV-2 RT-PCR (or any other diagnostic PCR for that matter) to counteract PCR dropouts due to mutations. With several targets a PCR drop out or relative shift in ct values compared to the other targets used can actually serve as an indication for a mutation, prompting further investigation of the sample by genome sequencing. Furthermore, it emphasizes the importance to remain vigilant with regard to new mutations and their potential effect on PCR performance. In this context it is also important to keep in mind the underrepresentation of genome data from some geographical regions (e.g. Africa), which may lead to mutations from these regions being overlooked, despite causing problems in diagnostics in these regions.

## 5. Acknowledgements

The work was funded by the Medical Service of the German Ministry of Defense.

## 6. Author contributions

Fee Zimmermann – conceptualization, methodology, formal analysis, visualization, writing: review and editing, supervision, Maria Urban – investigation, validation, writing: review and editing, Christian Krüger - investigation, Mathias Walter - software, writing: review and editing, Roman Wölfel – formal analysis, writing: review and editing, Katrin Zwirglmaier – conceptualization, methodology, formal analysis, writing: original draft, supervision

## Supplementary information

**Supplementary Table. 1.**
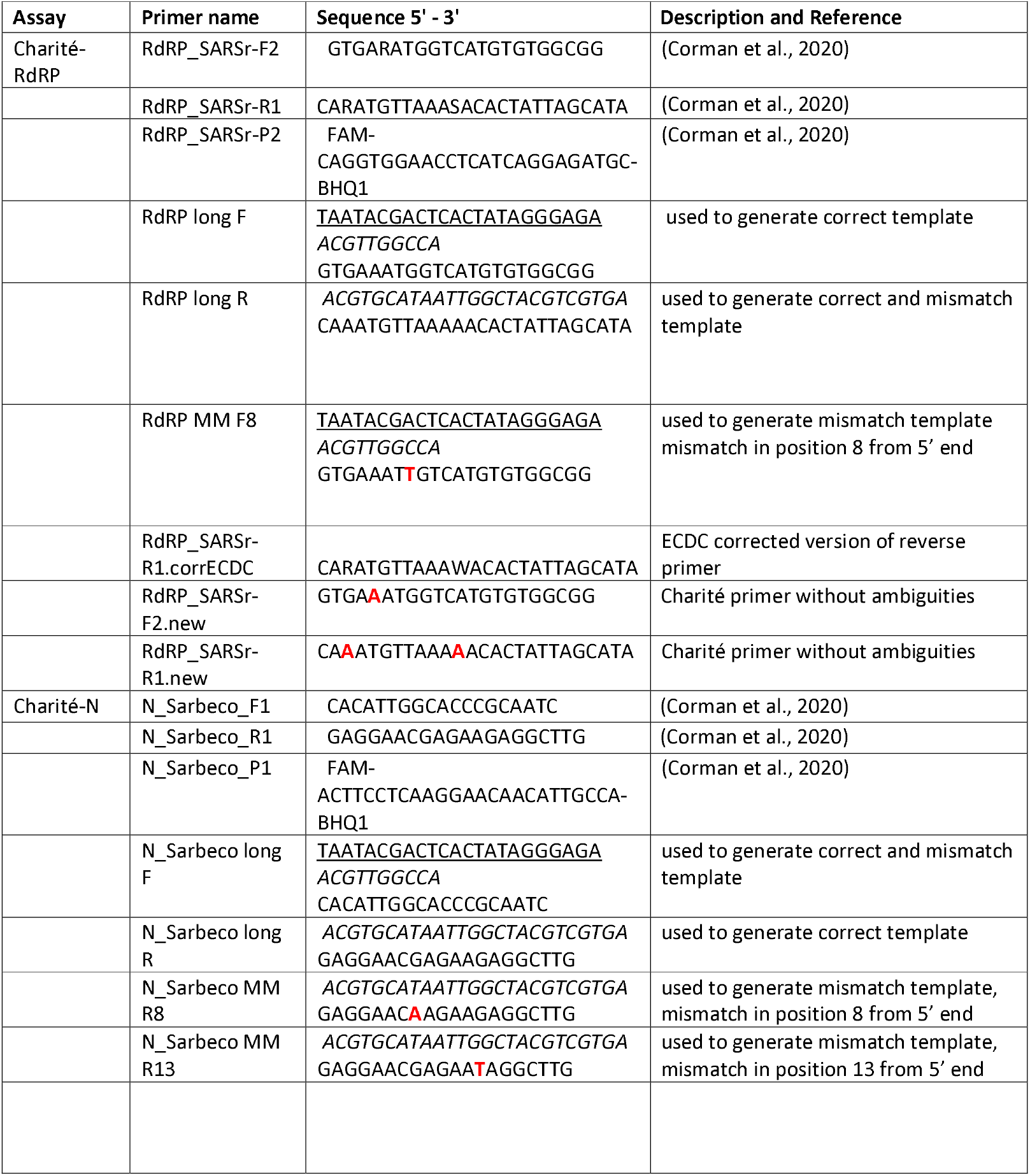
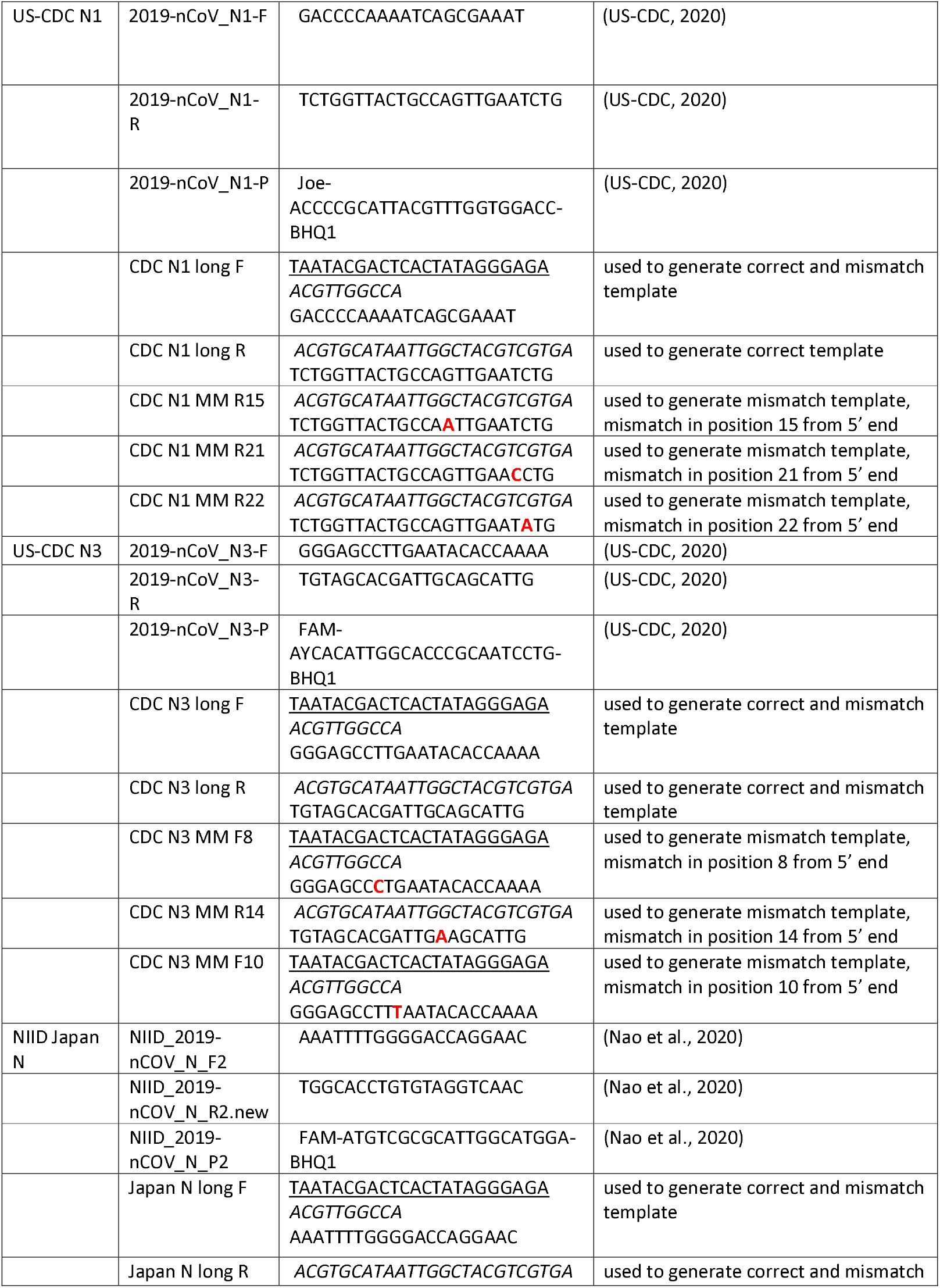
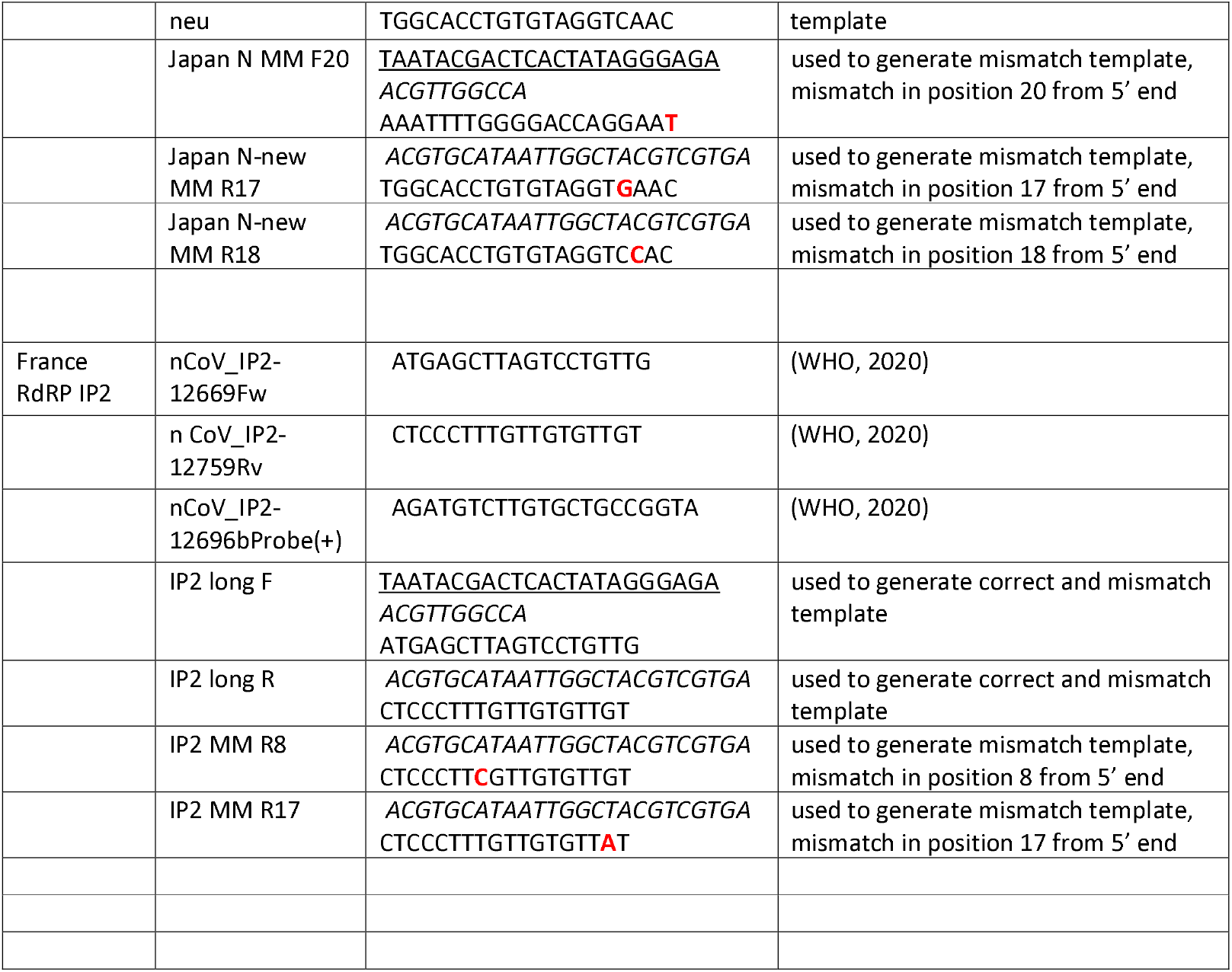
Primer and probe sequences used in this study. Underlined = T7 promotor, italicized = nonsense sequence to extend the amplicon, bold and red = mismatch

## Notes

### Competing Interest Statement

The authors have declared no competing interest.

